# The *GASA* Gene Family in *Theobroma cacao*: Genome wide Identification and Expression Analysis

**DOI:** 10.1101/2021.01.27.425041

**Authors:** Abdullah, Sahar Faraji, Furrukh Mehmood, Hafiz Muhammad Talha Malik, Ibrar Ahmed, Parviz Heidari, Peter Poczai

**Author notes:** Correspondence: Abdullah, Parviz Heidari, Péter Poczai.

## Abstract

The gibberellic acid-stimulated *Arabidopsis* (*GASA/GAST*) gene family is widely distributed in plants. The role of the *GASA* gene family has been reported previously in various physiological and biological processes, such as cell division, root and seed development, stem growth, and fruit ripening. These genes also provide resistance to abiotic and biotic stresses including antimicrobial, antiviral, and antifungal. Here, we report 17 *tcGASA* genes in *Theobroma cacao* L. distributed on six chromosomes. The gene structure, promoter-region sequences, protein structure, and biochemical properties, expression, and phylogenetics of all *tcGASA*s were analyzed. Phylogenetic analyses divided tcGASA proteins into five groups. The nine segmentally duplicating genes form four pairs and cluster together in phylogenetic tree. Purifying selection pressure was recorded on *tcGASA*, including duplicated genes. Several stress/hormone-responsive cis-regulatory elements were also recognized in the promoter region of *tcGASAs.* Differential expression analyses revealed that most of the *tcGASA* genes showed elevated expression in the seeds (cacao food), implying their role in seed development. The black rod disease of genus *Phytophthora* caused up to 20–25% loss (700,000 metric tons) in world cacao production. The role of *tcGASA* genes in conferring fungal resistance was also explored based on RNAseq data against *Phytophthora megakarya*. The differential expression of *tcGASA* genes was recorded between the tolerant and susceptible cultivars of cacao plants, which were inoculated with the fungus for 24h and 72h. This differential expression indicating possible role of *tcGASA* genes to fungal resistant in cacao. Our findings provide new insight into the function, evolution, and regulatory system of the *GASA* family genes in *T.* cacao and provide new target genes for development of fungi-resistant cacao varieties in breeding programs.

## Introduction

The gibberellic acid–stimulated *Arabidopsis* (*GASA/GAST*) gene family is widely distributed in plants and performs various functions (Ahmad et al., 2020b; Fan et al., 2017). *GAST1* was the first gene identified among the *GASA* family’s genes in tomato (Shi et al., 1992). GASA proteins are comprised of three domains, including a signal peptide of up to 18–29 amino acids at the N-terminal, hydrophilic and high variable regions of up to 7–31 amino acids in the center, and a conserved domain at the C-terminal of up to 60 amino acids which mostly includes 12 cysteine residues (Rezaee et al., 2020; Silverstein et al., 2007; Zhang and Wang, 2008). This third domain of the C-terminal is the characteristics of all identified GASAs (Ahmad et al., 2019; Fan et al., 2017; Rezaee et al., 2020; Zhang and Wang, 2008). These cysteine-rich peptides play a vital role in various plant processes. Their roles have been stated in organ development (Fuente et al., 2006), lateral root development (Zimmermann et al., 2010), stem growth (Zhang et al., 2009), cell division (Nahirñak et al., 2012; Roxrud et al., 2007), fruit ripening and development (Moyano-Cañete et al., 2013), flowering time (Ahmad et al., 2019; Fan et al., 2017; Zhang et al., 2009), seed development (Ahmad et al., 2020b; Roxrud et al., 2007), and bud dormancy (Yang et al., 2019). The detailed expression analyses of various tissues of Arabidopsis, tomato, rice, soybean, and apple showed the tissue-specific expression of the *GASA* genes (Ahmad et al., 2019; Fan et al., 2017; Rezaee et al., 2020). For example, *GASA* genes in tomato, including *Solyc11g011210*, *Solyc12g089300*, and *Solyc01g111075,* showed high expression at the fruit-ripening stage, whereas *Solyc03g113910*, *Solyc01g111075*, *Solyc06g069790*, and *Solyc12g042500* showed high expression level at the flowering stage (Rezaee et al., 2020). Moreover, some studies also showed the contrast effect of GASA proteins. For instance, *AtGASA4* promotes flowering (Roxrud et al., 2007), while *AtGASA5* induces the opposite effect (Zhang et al., 2009).

The *GASA* genes also play crucial roles in response to various biotic, abiotic, and hormone-related stresses. For instance, a member of the *GASA* gene family, *GmSN1*, provides resistance to viruses in soybean and Arabidopsis (He et al., 2017). Similarly, a high expression level of *CcGASA4* was reported in citrus leaves after infection with *Citrus tristeza* virus (Wu et al., 2020). The antimicrobial properties of various proteins of the GASA family have also been reported (Almasia et al., 2008; Berrocal-Lobo et al., 2002; Kovalskaya and Hammond, 2009). The antifungal activity of the GASA proteins has been reported in almost all tissues of potato, including root, tubers, leaves, stem, stolon, axillary bud, and flowers (Almasia et al., 2008; Berrocal-Lobo et al., 2002; Kovalskaya and Hammond, 2009). Similarly, the antifungal activity of GASA members has been found in Arabidopsis, tomato, Alfalfa, and Jujuba (Balaji and Smart, 2012; García et al., 2014; Qu et al., 2016; Zhang and Wang, 2008). The GASAs also showed resistance to various abiotic stresses, such as salt and drought (Wang et al., 2018). The induction of *GASA4* and *GASA6* has been reported in Arabidopsis by growth hormones such as auxin, BR, GA, and cytokinin. In contrast, repression has been stated by stress hormones including SA, ABA, and JA (Qu et al., 2016).

*Theobroma cacao* L. belongs to the family Malvaceae (Purseglove, 1968). This is an economically important tree and grows in up to 50 countries located in the humid tropics (Motamayor et al., 2013). *Theobroma cacao* L. seeds are enclosed in pods and are used for chocolate production, confectionery, and cosmetics (Litz, 2005). This plant is adapted to high humidity areas, and is therefore predisposed to various fungal diseases (Bridgemohan and Mohammed, 2019; McElroy et al., 2018). Pod rot, or black rod, is caused by *Phytophthora* species of fungus (*P. megakarya*, *P. palmivora*, and *P. capsici*), leading to 20–30% loss in yield and 10% death of trees (Bridgemohan and Mohammed, 2019). The elucidation of the whole genome is helping to understand the genetic bases of biotic and abiotic stresses (Motamayor et al., 2013). The availability of the high-quality chromosome-level genome assembly of *Theobroma cacao* (Argout et al., 2017; Motamayor et al., 2013) provides quality resources for the characterization of various gene families to elucidate the role of different gene families in cacao development. However, few gene families such as WRKY (Dayanne et al., 2017) and NAC (Shen et al., 2020) have been elucidated in *Theobroma cacao*. Moreover, genome-wide analyses were also performed to reveal the divergent patterns of gene expression during zygotic and somatic embryo maturation in cacao (Maximova et al., 2014). Here, we are interested in characterizing the *GASA* gene family and providing data about the comprehensive and expression analyses of this gene family specifically regarding fungus-related diseases.

The expression pattern and evolutionary relationships of *GASA* genes were studied in Arabidopsis (Zhang and Wang, 2008), apple (Fan et al., 2017), common wheat (Cheng et al., 2019), grapevine (Ahmad et al., 2020b), soybean (Ahmad et al., 2019), and *Solanum tuberosum* subsp. tuberosum cv. Kennebec (Nahirñak et al., 2016). In the current study, we are interested in identifying and characterizing GASAs of *T. cacao* and exploring their roles in various biotic and abiotic stresses. We provide data of GASAs related to their distribution in genome, chemical properties, subcellular localization, and cis-regulatory elements of promoter regions for the first time in cacao. We also explored the roles of GASA in various abiotic and biotic stresses. This helps us to identify GASAs that show high expressions against infections of fungus *Phytophthora megakarya*.

## 2. Materials and Methods

### 2.1 Identification of *GASA* genes in the genome of *Theobroma cacao* and analyses for conserved GASA domain

We retrieved the GASA family’s protein sequences from Arabidopsis Information Resource (TAIR10) database (ftp://ftp.arabidopsis.org). We used them as a query in BLAST for the identification of *GASA* genes in the *Theobroma cacao* genome. The *GASA* genes were identified in the latest version of the *Theobroma cacao* genome (*Theobroma cacao* Belizian Criollo B97-61/B2) (Argout et al., 2017). We retrieved protein sequences, coding DNA sequences (CDS), genomics, and promoter sequence (1500 bp upstream of gene). The retrieved protein sequences were further analyzed for the presence of the GASA domain using CDD (https://www.ncbi.nlm.nih.gov/Structure/cdd/wrpsb.cgi) database of the National Center for Biotechnology Information (NCBI). All the sequences that showed the presence of the conserved GASA domain were selected for further analyses, whereas all the proteins with an absent/truncated domain were discarded.

### 2.2 Chromosome mapping and characterization of physiochemical properties

The position of each gene, including chromosome number and position on chromosome, was noted. All the genes were renamed according to the location of the chromosome and their position, as shown in table 1. The MapChart software (Voorrips, 2002) through Ensemble was used to show the position of each *tcGASA* gene along the position of the chromosome. Various physiochemical properties, including length of the protein, molecular weight (MW), isoelectric point (PI), instability index, and the grand average of hydropathy (GRAVY), were determined using the ExPASy tool (Gasteiger et al., 2005). The subcellular localization of *GASA* genes was also predicted using the BUSCA webserver (Savojardo et al., 2018).

**Table 1.** List of the identified *GASA* genes and their characteristics in *Theobroma cacao*.

| Gene locus ID | Gene ID | Location | Length(aa) | MW (kDa) | pI | Instability index | GRAVY | Subcellular localization |
| --- | --- | --- | --- | --- | --- | --- | --- | --- |
| Tc01v2_t001590 | *TcGASA1* | 1: 830,757-831,466 | 105 | 11.18 | 8.93 | Unstable | -0.023 | Extracellular space |
| Tc01v2_t005890 | *TcGASA2* | 1: 3,268,900-3,269,687 | 107 | 11.44 | 8.98 | Unstable | -0.057 | Extracellular space |
| Tc02v2_t009150 | *TcGASA3* | 2: 5,635,674-5,636,363 | 90 | 9.90 | 9.01 | Stable | -0.054 | Extracellular space |
| Tc02v2_t021110 | *TcGASA4* | 2: 30,717,916-30,719,013 | 106 | 11.91 | 9.21 | Unstable | -0.195 | Extracellular space |
| Tc04v2_t000240 | *TcGASA5* | 4: 270,172-272,304 | 320 | 33.92 | 9.83 | Unstable | -0.750 | Extracellular space |
| Tc04v2_t006890 | *TcGASA6* | 4: 15,324,400-15,325,648 | 106 | 11.83 | 9.32 | Unstable | -0.343 | Extracellular space |
| Tc04v2_t009440 | *TcGASA7* | 4: 19,321,294-19,322,037 | 95 | 10.37 | 9.26 | Unstable | -0.072 | Extracellular space |
| Tc04v2_t016520 | *TcGASA8* | 4: 26,297,870-26,298,743 | 123 | 13.42 | 8.27 | Unstable | -0.026 | Plasma membrane |
| Tc04v2_t016530 | *TcGASA9* | 4: 26,301,782-26,302,647 | 95 | 10.62 | 8.88 | Unstable | -0.251 | Extracellular space |
| Tc05v2_t012230 | *TcGASA10* | 5: 25,218,365-25,220,141 | 114 | 12.76 | 9.46 | Unstable | -0.327 | Extracellular space |
| Tc08v2_t003650 | *TcGASA11* | 8: 1,941,175-1,942,036 | 102 | 10.92 | 6.67 | Stable | -0.031 | Extracellular space |
| Tc08v2_t003660 | *TcGASA12* | 8: 1,943,144-1,944,201 | 100 | 10.57 | 8.66 | Unstable | 0.004 | Extracellular space |
| Tc08v2_t007890 | *TcGASA13* | 8: 4,555,655-4,556,418 | 88 | 9.64 | 9.32 | Stable | -0.062 | Extracellular space |
| Tc08v2_t014670 | *TcGASA14* | 8: 15,650,553-15,651,438 | 102 | 10.98 | 8.72 | Unstable | -0.164 | Extracellular space |
| Tc08v2_t014680 | *TcGASA15* | 8: 15,651,664-15,652,892 | 92 | 10.01 | 8.90 | Stable | -0.148 | Extracellular space |
| Tc08v2_t014690 | *TcGASA16* | 8: 15,653,408-15,654,378 | 110 | 12.13 | 9.64 | Unstable | -0.263 | Extracellular space |
| Tc09v2_t020100 | *TcGASA17* | 9: 30,243,952-30,245,432 | 112 | 12.32 | 9.46 | Unstable | -0.252 | Extracellular space |

### 2.3 Gene structure and promoter region analyses

We analyzed CDS sequences for exons–introns within all *tcGASA* genes using the Gene Structure Display Server (http://gsds.cbi.pku.edu.cn). PlantCare (Lescot et al., 2002) was used to study cis-regulatory elements in the 1500 bp promoter region.

### 2.4 Prediction of post-translational modifications of GASA proteins

The phosphorylation site of the GASA proteins was predicted by the NetPhos 3.1 server (http://www.cbs.dtu.dk/services/NetPhos/) (Blom et al., 2004) with a potential value > 0.5. N-glycosylation sites were predicted using the NetNGlyc 1.0 server (http://www.cbs.dtu.dk/services/NetNGlyc/) (Gupta and Brunak, 2002) with default parameters.

### 2.5 Phylogenetic and conserved motif analyses

The phylogenetic relationship of the *GASA* genes of *T. cacao* was inferred with *GASA* genes of six other species, including Arabidopsis, *Gossypium raimondii*, *Vitis vinifera*, rice, *Brachypodium distachyon*, and maize. A similar approach, which was given for the identification of *GASA* genes in *Theobroma cacao,* was employed for the identification of the genes of all other species. Clustal Omega (https://www.ebi.ac.uk/Tools/msa/clustalo/) (Sievers et al., 2011) was used for multiple alignment of the protein sequences of all species. The unrooted neighbor-joining tree was drawn using MEGA X (Kumar et al., 2018) and visualization of the tree was improved by using an interactive tree of life (iTOL) (Letunic and Bork, 2019). The conserved motif distribution into GASA proteins was performed using MEME v5.3.0 server (http://meme-suite.org/tools/meme) (Bailey et al., 2009). We searched for a maximum number of five motifs with a minimum width of motif 6 and a maximum motif width of 30.

### 2.6. Gene duplications and estimation of Ka/Ks values

The identity of > 85% in nucleotide sequences of genes is considered a sign of duplication (Zheng et al., 2010). Hence, we aligned DNA coding sequences using Clustal Omega (Sievers et al., 2011), and the extent of the identity of the genes with each other was determined using Geneious R8.1 (Kearse et al., 2012). Gene duplication events, as compared to other species, were determined using the MCScan v0.8 program (Wang et al., 2012) through the Plant Genome Duplication Database.

The selection pressure on on *GASA* genes was determined by calculating non-synonymous (Ka) and synonymous (Ks) substitutions and the ratio of non-synonymous to synonymous substitutions (Ka/Ks) using DnaSP v6 software (Rozas et al., 2017). The time divergence and duplication were assessed by a synonymous mutation rate of λ substitutions per synonymous site per year as T = (Ks/2λ) (λ = 6.5 × 10^−9^) × 10^−6^ (Yang et al., 2008). The synteny relationships of *GASA* genes among the orthologous pairs of cacao−rice, cacao-Arabidopsis, cacao-maize, cacao–*Brachypodium distachyon*, cacao-*Gossypium raimondii*, and cacao−*Vitis vinifera* at both gene and chromosome levels were visualized using Circos software (Krzywinski et al., 2009).

### 2.7 Three-dimensional protein modeling and molecular docking

We used iterative template-based fragment assembly simulations in I-TASSER (Yang et al., 2015) to build three-dimensional protein structures of GASAs after selection of best models by the 3D-refine program (Bhattacharya et al., 2016). We draw a Ramachandran plot using the RAMPAGE program to validate the predicted structures by measuring backbone psi (ψ) and dihedral phi (◻) angles (Lovell et al., 2003). We also used P2Rank in PrankWeb software (Jendele et al., 2019) and CASTp tool (Tian et al., 2018) to analyze the refined structure of GASA proteins to predict protein pockets and cavities. Finally, PyMOL (DeLano, 2002) was used to visualize results.

### 2.8. In silico expression analysis of *GASA* Genes through RNA-seq data

The publicly available RNA-seq data related to the cacao genome was employed for expression assays to measure *GASA* family members in multiple tissues and during various biotic and abiotic stimuli exposure. The RNA-seq data of cacao under various time courses of biotic stress (*Phytophthora megakarya*) were downloaded from GEO DataSets under accession number GSE116041 (Pokou et al., 2019). This data was also log2 transformed to generate heatmaps via the TBtools package (Chen et al., 2020). Furthermore, the expression levels of *GASA* genes for tissue specific expression and in under multiple abiotic stresses, including cold, osmotic, salt, drought, UV, wounding and heat, have been detected in the *Arabidopsis* orthologous genes for *tcGASAs* (SAMEA5755003 and PRJEB33339).

## 3. Results

### 3.1 Identification of *GASA* genes and their distributions on chromosomes within genomes

We detected 17 *GASA* genes in the genome of *T. cacao* distributed on six chromosomes out of ten. These genes were named from *tcGASA1* to *tcGASA17* based on their distribution on chromosomes starting from chromosome 1. When two or more genes were present on the same chromosome, then the gene present at the start of a chromosome was named first (Table 1 and Figure 1). Six genes were distributed on chromosome 8 and five genes were distributed on chromosome 4. Chromosome 1 and chromosome 2, each contained two genes, whereas chromosome 5 and chromosome 9 contained one gene each. This data showed the unequal distribution of *tcGASA* genes within the cacao genome. The location of each gene on the chromosome is mentioned in Table 1, as well as the start and end. Moreover, the sequences of genes, proteins, coding regions, and promoter regions are provided in Table S1.

**Figure 1.**
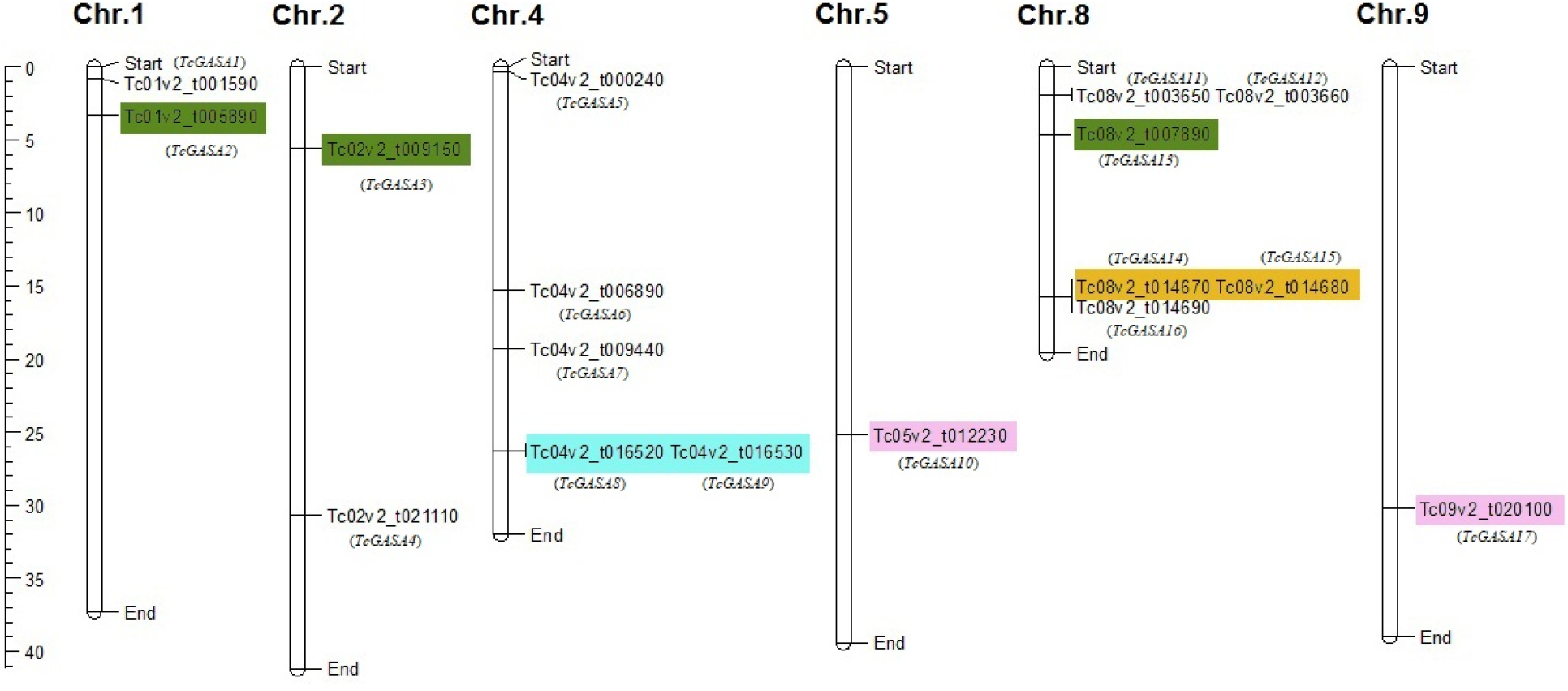
Location of *tcGASA* genes on the chromosome in *Theobroma cacao*. Each gene is shown with a gene identification number along with the number given to each gene in the current study. The pairs of segmentally duplicated genes are shown with same color.

### 3.2 Protein length, molecular weight, and isoelectric point of tcGASA proteins

In the current study, tcGASAs were characterized based on their physiochemical properties (Table 1). The identified tcGASA proteins were low molecular weight proteins ranging in length from 88 (tcGASA13) to 320 (tcGASA5) amino acids with molecular weight (MW) ranging from 9.64 to 33.92 kDa. Except for tcGASA5 and tcGASA8, with molecular weights of 13.42 kDa and 33.92 kDa, respectively, the MW of all other tcGASAs was found to be less than 13 kDa. The isoelectric point also showed similarities among tcGASA and suggested the alkaline nature of the proteins. Except for tcGASA11, which has an isoelectric point of 6.67, all other tcGASA proteins have isoelectric points of more than 8, ranging from 8.27 to 9.83.

### 3.3 Analyses of instability index, GRAVY, and subcellular localization of tcGASA proteins

The instability index provides information about the stable and non-stable features of proteins in the various biochemical processes. The instability index indicated 4 stable tcGASAs including tcGASA3, tcGASA11, tcGASA13, and tcGASA15, as well as 13 unstable tcGASAs (Table 1). The positive value of GRAVY indicates its hydrophobic nature, whereas the negative value indicates the hydrophilic nature of proteins. The GRAVY value was recorded as negative for sixteen tcGASA proteins ranging from −0.75 to −0.023, but as positive (0.004) for tcGASA12 (Table 1). Hence, the data indicate the hydrophilic nature of most tcGASA proteins. Subcellular localization provides information about the function of proteins. Based on BUSCA, we predicted extracellular localization of tcGASA proteins, except for tcGASA8, which localized in the plasma membrane (Table 1).

### 3.4 tcGASA proteins 3D structure analyses and post-translational modifications

The predicted 3D structure of all tcGASA proteins showed that these proteins contain β sheets, α helices, random coils, and extended strands. The random coils were the most abundant and were more extensive than α helices, while the β sheets were the least (Figure 2). In the present study, the post-translational modifications of tcGASAs were predicted in terms of phosphorylation and glycosylation (Figure 3, Table S2). We predicted a total of 224 potential phosphorylation events on amino acids serine, threonine, and tyrosine within tcGASA proteins. Most of the phosphorylation events were predicted related to serine (92) followed by threonine (86) and then by tyrosine (46). Among tcGASA proteins, most of phosphorylation sites (57 sites) were predicted in tcGASA05, whereas in other proteins’ phosphorylation events ranged from 9 to 14 sites. Three tcGASA including tcGASA10, tcGASA15, and tcGASA17, were also identified with a potential glycosylation site (Table S2).

**Figure 2.**
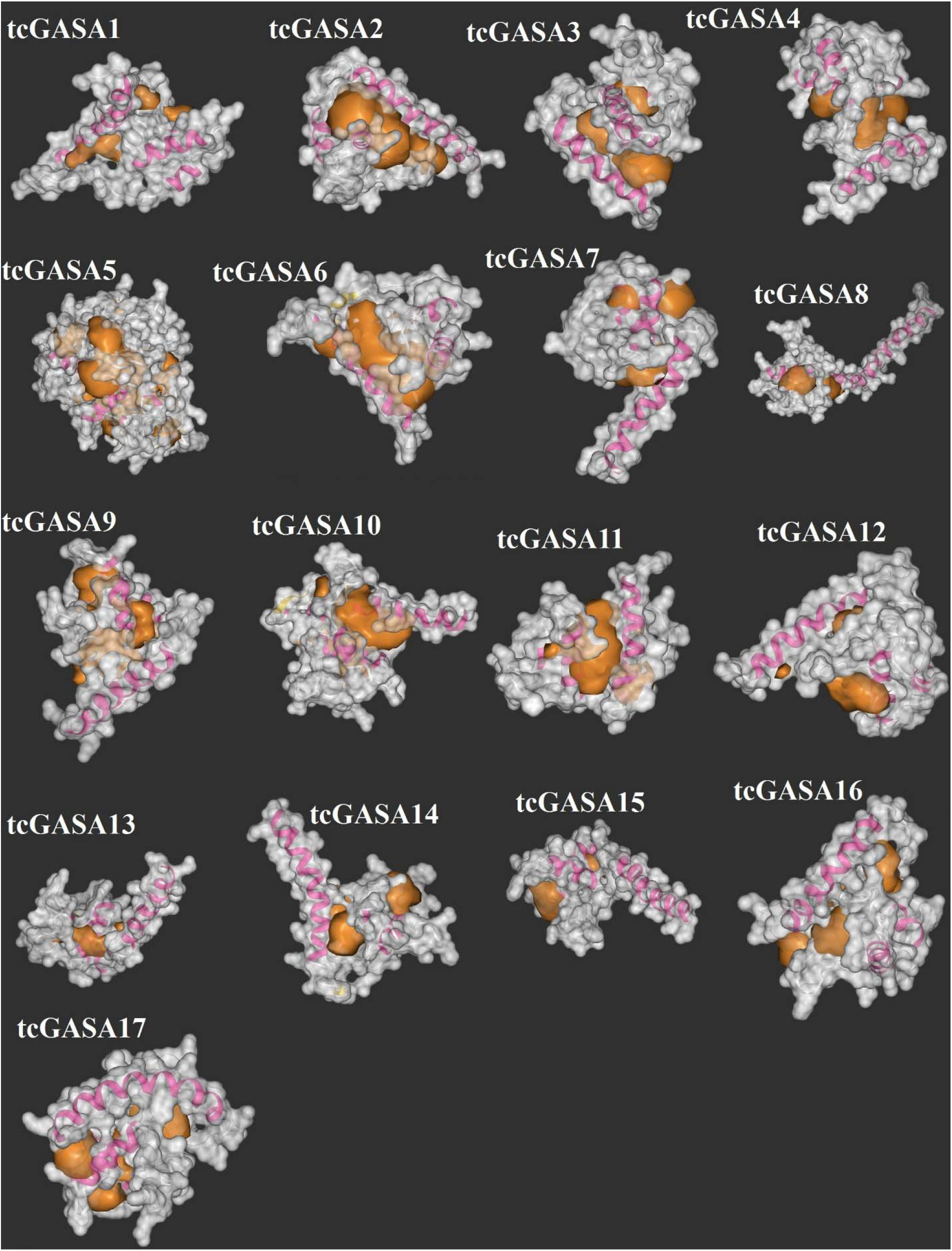
3D structure analyses of tcGASA proteins.

**Figure 3.**
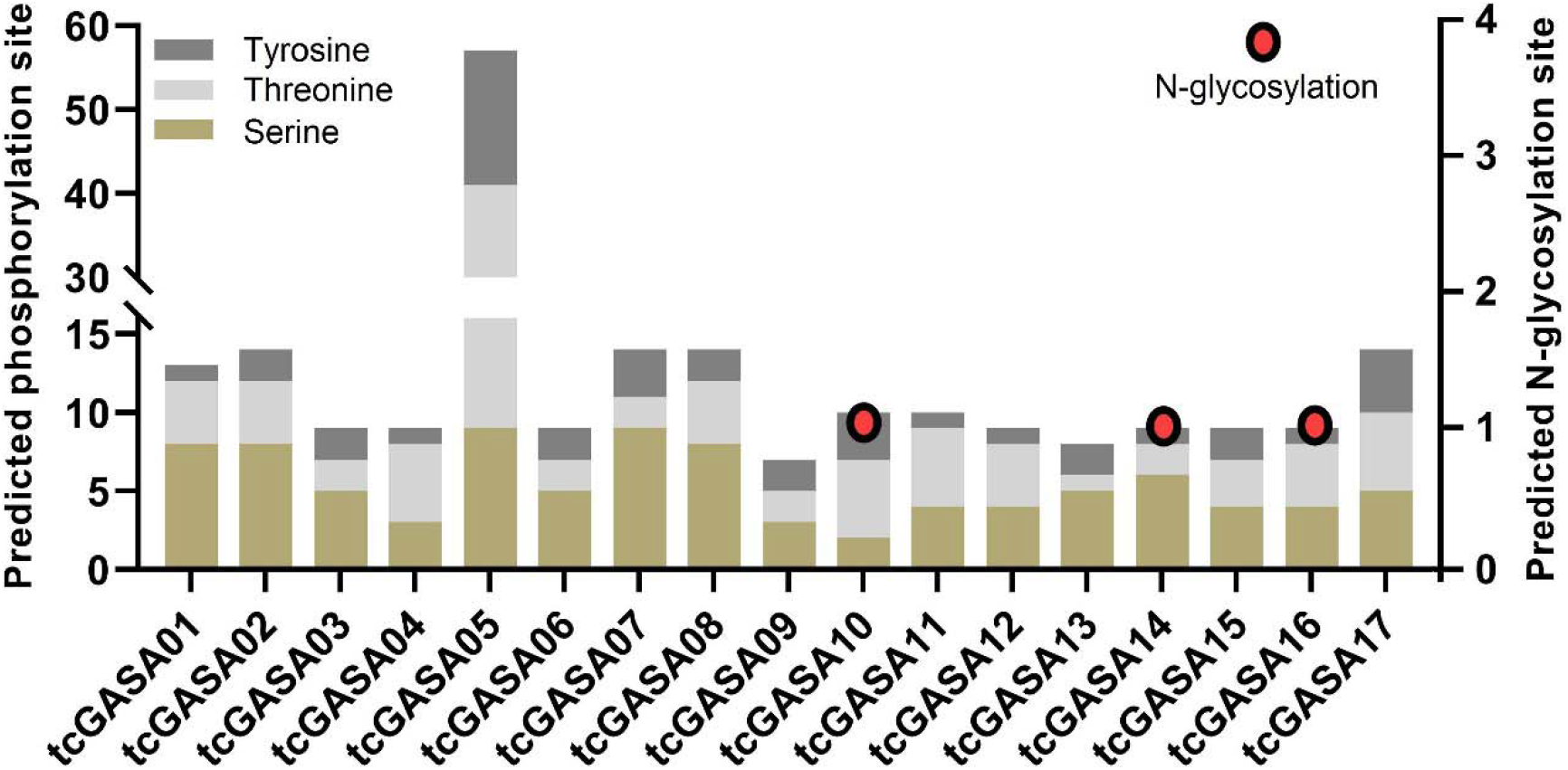
Post translational phosphorylation and glycosylation modification of tcGASA proteins. The circle is drawn on proteins in which the glycosylation site was predicted.

### 3.5 Phylogenetic analyses of tcGASA proteins

The phylogenetic inference of ninety-nine GASA-protein sequences from seven species (mentioned in methodology), including 17 sequences of tcGASA, resolved into five groups (Figure 4). The GASA proteins of cacao showed high variations and were distributed into all five groups. Group I include only one sequence tcGASA16 (Tc08v2_t014690). Group II was confined to five protein sequences including tcGASA01 (Gene ID: Tc01v2_t001590), tcGASA4 (Tc02v2_t021110), tcGASA5 (Tc04v2_t000240), tcGASA11 (Tc08v2_t003650), and tcGASA12 (Tc08v2_t003660). Group III comprised two sequences—tcGASA14 (Tc08v2_t014670) and tcGASA15 (Tc08v2_t014680). Group IV included tcGASA2 (Tc01v2_t005890), tcGASA03 (Tc02v2_t009150), and tcGASA13 (Tc08v2_t007890). Group V consisted of six protein sequences, including tcGASA6 (Tc04v2_t006890), tcGASA7 (Tc04v2_t009440), tcGASA8 (Tc04v2_t016520), tcGASA9 (Tc04v2_t016530), tcGASA10 (Tc05v2_t012230), and tcGASA17 (Tc09v2_t020100). The tcGASA4 (Tc02v2_t021110) shared a node with one of the protein sequences of *Vitis vinifera,* whereas the other 16 proteins showed close relationships with the sequences of another species of family Malvaceae (*Gossypium raimondii*).

**Figure 4.**
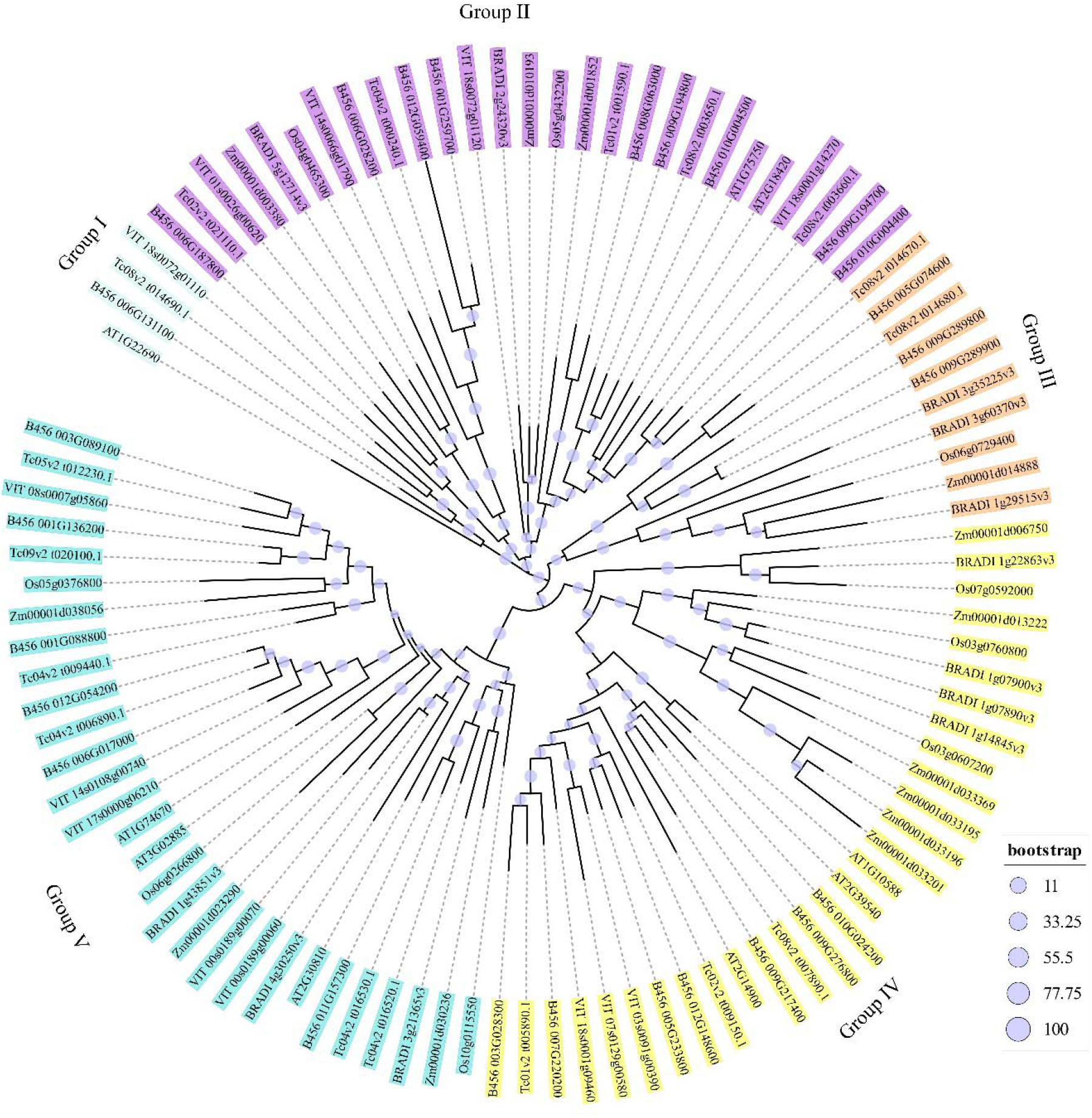
The phylogenetic inference of tcGASA proteins along with protein sequences of six other species. Each sequence is presented with gene number. The start of each sequence contained code for the species as: Tc: *Theobroma cacao*; AT: *Arabidopsis thaliana*; B456: *Gossypium raimondii*; VIT: *Vitis vinifera*; Os: *Oryza sativa* (rice); BRADI: *Brachypodium distachyon*; Zm: *Zea mays* (Maize).

### 3.6 Gain and loss of intron(s), proteins motifs, and their linked to phylogenetic grouping

We also determined numbers of introns-exons within the genes and motifs in protein sequences. We drew a separate phylogeny of the *tcGASA* genes to compare *tcGASA* genes for presence and location of introns, and tcGASA proteins for conserved motifs, based on phylogenetic clusters that they formed. Each phylogenetic group was shown with different colors for clarity (Figure 5.a). The analyses of genomic sequences showed the absence of intron in one gene, presence of one intron in five genes, two introns in eleven genes, and three introns in one gene (Figure 5b). Genes clustered together showed differences in their number and in intron-exon distributions (Figure 5a, b). Five motifs were revealed in tcGASA. Four motifs (1–4) were distributed in all tcGASAs, whereas a fifth motif was limited to Tc04v2_t016520 (tcGASA8), Tc05v2_t012230 (tcGASA10), and Tc09v2_t020100 (tcGASA17) (Figure 5c). The motifs of proteins that clustered together within the phylogenetic tree presented similarities in the distribution of motifs to some extent (Figure 5 a, c).

**Figure 5.**
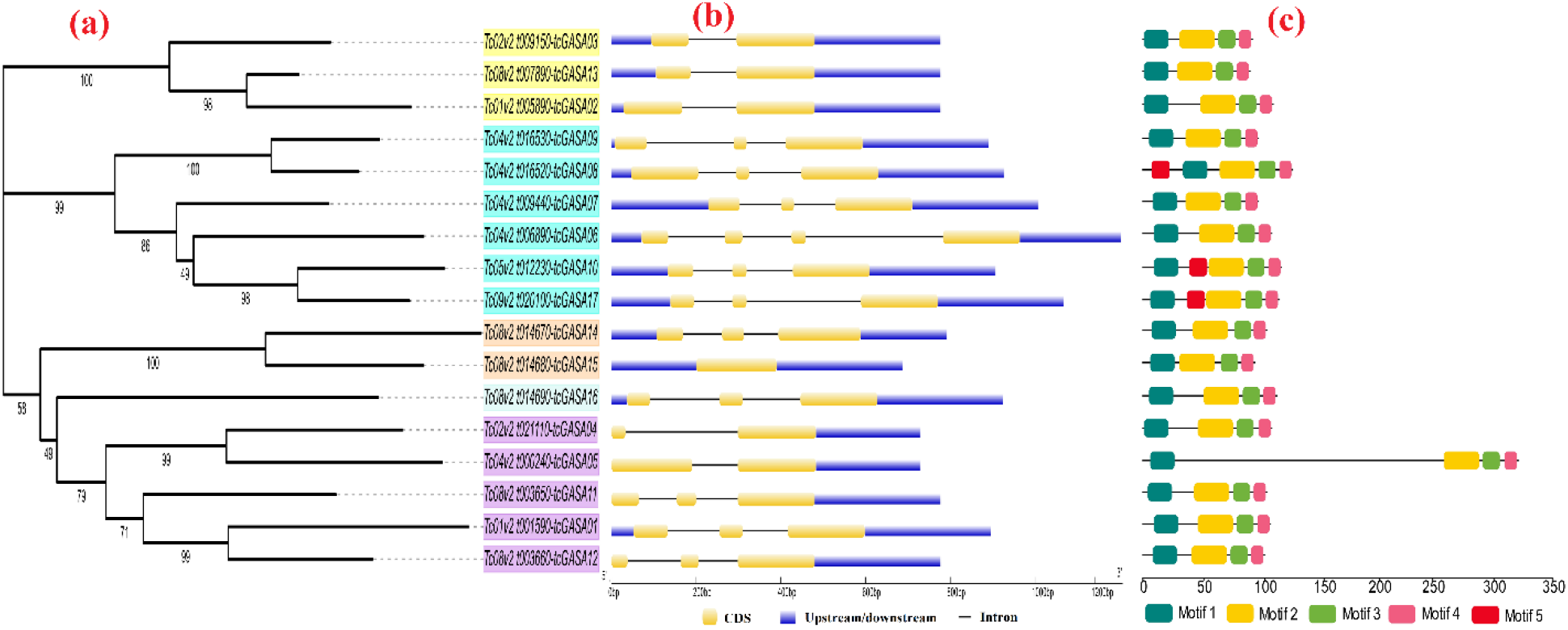
The phylogenetic analyses in comparison to introns-exons in genes and motifs in proteins of tcGASAs. (a) Phylogeny of tcGASA, (b) introns-exons in *tcGASA* genes, (c) Motifs in tcGASA proteins.

### 3.7 Duplications, divergence, and synteny among GASA genes

We analyzed the paralogous relationships among the *GASA* genes within cacao and analyzed the orthologous relationships of *GASA* genes by comparing them with the other six species (mentioned in methodology). Segmental duplications were found in nine *GASA* genes that were paired into four groups as: *tcGASA*2-*tcGASA*3-*tcGASA*13; *tcGASA*8-*tcGASA*9; *tcGASA*10-*tcGASA*17; and *tcGASA*14-*tcGASA*15 (Table 2) (as shown with similar colors in Figure 1). The analyses of synonymous and non-synonymous substitutions revealed that high purifying selection pressure exists on these genes after duplication. The analyses of divergence time indicated that the event of duplication of these pairs occurred 50 MYA to 204 MYA (Table 2). The synteny analyses with *GASAs* of other species showed high resemblance and identified orthologous genes among *T. cacao* and compared species (Figure 6).

**Table 2.** Gene duplications, synonymous and non-synonymous substitutions and time of divergence.

| **Gene 1** | **Gene 2** | **Duplication type** | **K<sub>a</sub>** | **K<sub>s</sub>** | **K<sub>a</sub>/K<sub>s</sub>** | ***p*-value** | **Divergence time (MYA)** |
| --- | --- | --- | --- | --- | --- | --- | --- |
| *TcGASA2* | *TcGASA3* | Segmental | 0.2679 | 2.6607 | 0.1007 | 1.04E-06 | 204.67 |
| *TcGASA2* | *TcGASA13* | Segmental | 0.1708 | 2.2495 | 0.0759 | 2.52E-08 | 173.04 |
| *TcGASA8* | *TcGASA9* | Segmental | 0.1515 | 0.6514 | 0.2325 | 1.30E-06 | 50.11 |
| *TcGASA10* | *TcGASA17* | Segmental | 0.283 | 1.0283 | 0.2752 | 3.28E-05 | 79.1 |
| *TcGASA14* | *TcGASA15* | Segmental | 0.3234 | 0.7971 | 0.4058 | 1.46E-08 | 61.32 |

**Figure 6.**
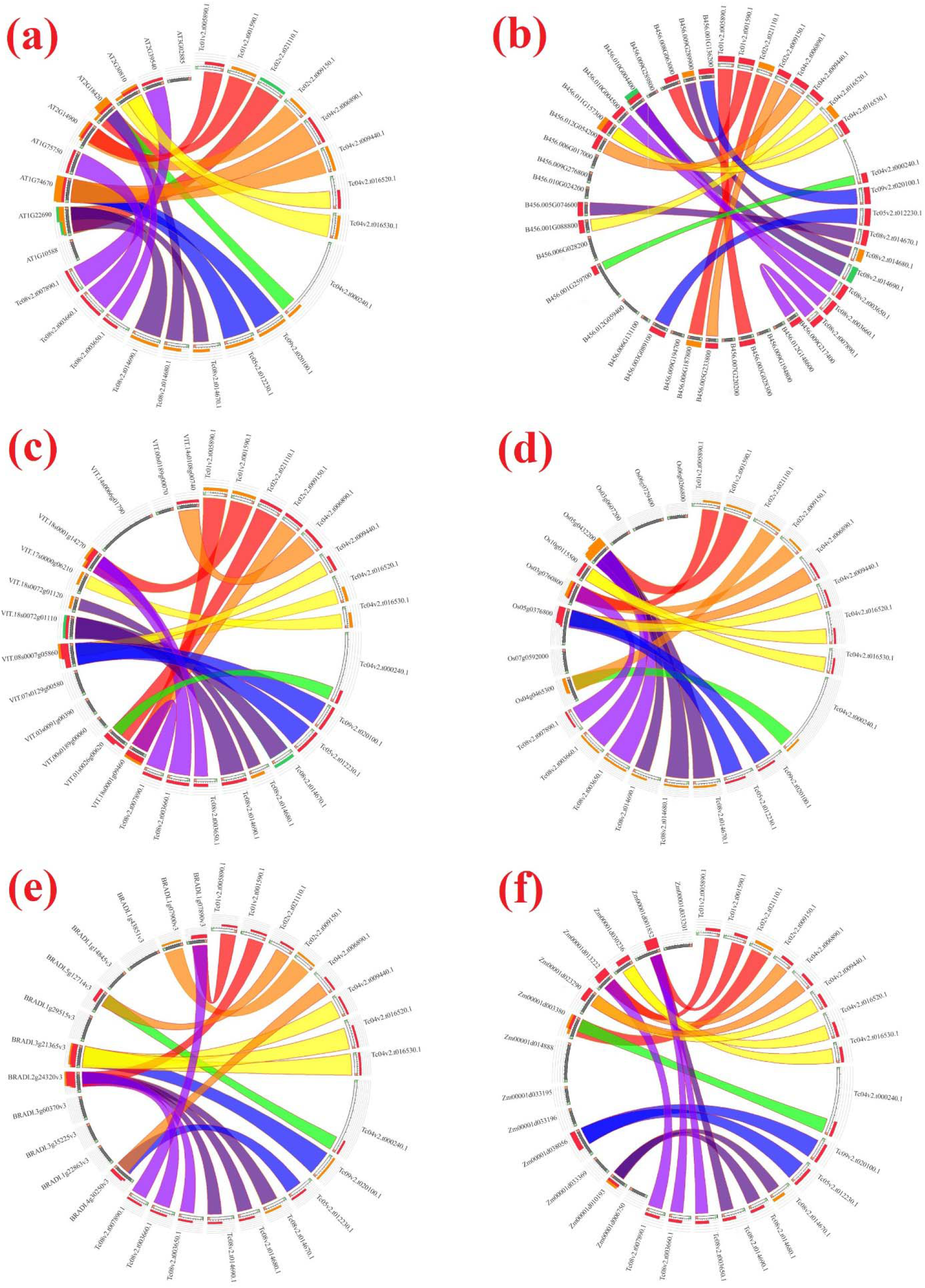
Synteny analysis of *GASA* genes. The syntenic blocks of cacao *GASA* genes are compared with six other species. (a) Arabidopsis, (b) *Gossypium raimondii*, (c) *Vitis vinifera*, (d) rice, (e) *Brachypodium distachyon*, and (f) maize.

### 3.8 Promoter regions analysis

The analyses of cis-regulatory elements in promoter regions revealed the presence of binding sites for key transcription factors related to light-responsive elements (48.88%), hormone-responsive elements (25.40%), stress-related elements (19.06%), growth-response elements (5.40%), and DNA- and protein-related binding sites (1.26) (Figure 7a). Regulatory sides were found for various hormones such as auxin, salicylic acid, abscisic acid, gibberellin, and MeJA (Figure 7b). Similarly, regulatory elements were identified for drought, elicitor, anaerobic induction, low temperature, and plant defense/stress (Figure 7c). The complete detail of each element, along with sequence and function is provided in Table S3.

**Figure 7.**
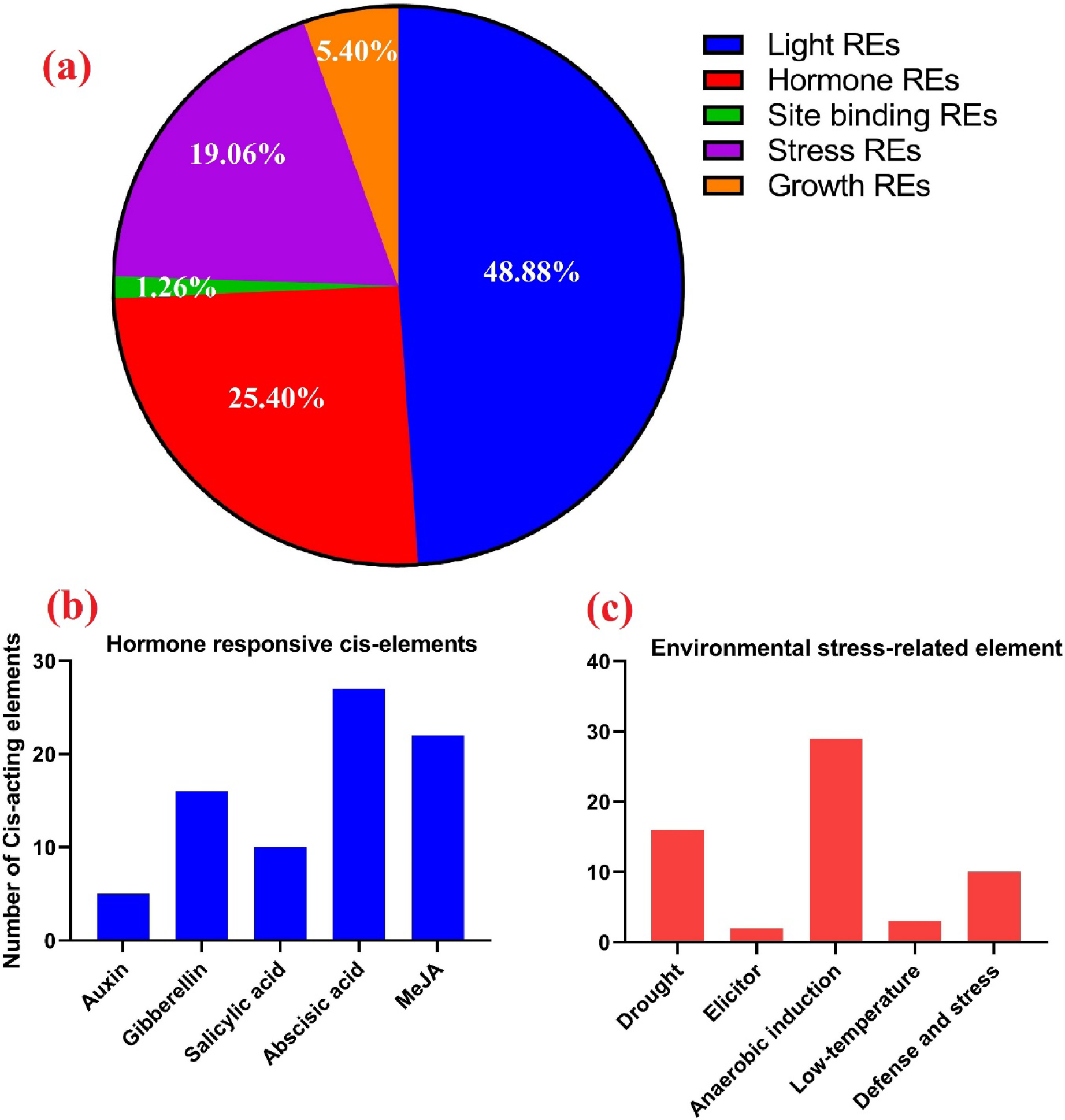
Cis-regulatory elements in promoter regions of *tcGASA* genes. (a) represents the extent of various types of regulatory elements based on function, such as light, hormone, growth, stress, and binding domain excluding the TATA box and CAAT box. (b) Distribution of different types of hormone-related cis-regulatory elements. (c) Distribution of cis-regulatory elements related to various types of environmental stresses.

### 3.9 Tissue-specific expression of *tcGASA* genes

We evaluated the expression of orthologous *tcGASAs* in various tissues to evaluate their role in the functions of *T. cacao* (Figure 8a). Differential expression was noted for *tcGASA* genes in various tissues of cacao. Five genes, including *tcGASA*02, *tcGASA*03, *tcGASA*08, *tcGASA*09, and *tcGASA*13, showed high expression in the beans (food part of cacao). In addition, *tcGASA*16 showed a high expression in leaves and entire seedlings. However, *tcGASA*02 and *tcGASA*03 were significantly downregulated in leaves compared to beans. *In-silico* expression results showed that *tcGASA*12 and *tcGASA*17 were less induced in pistil tissues.

**Figure 8.**
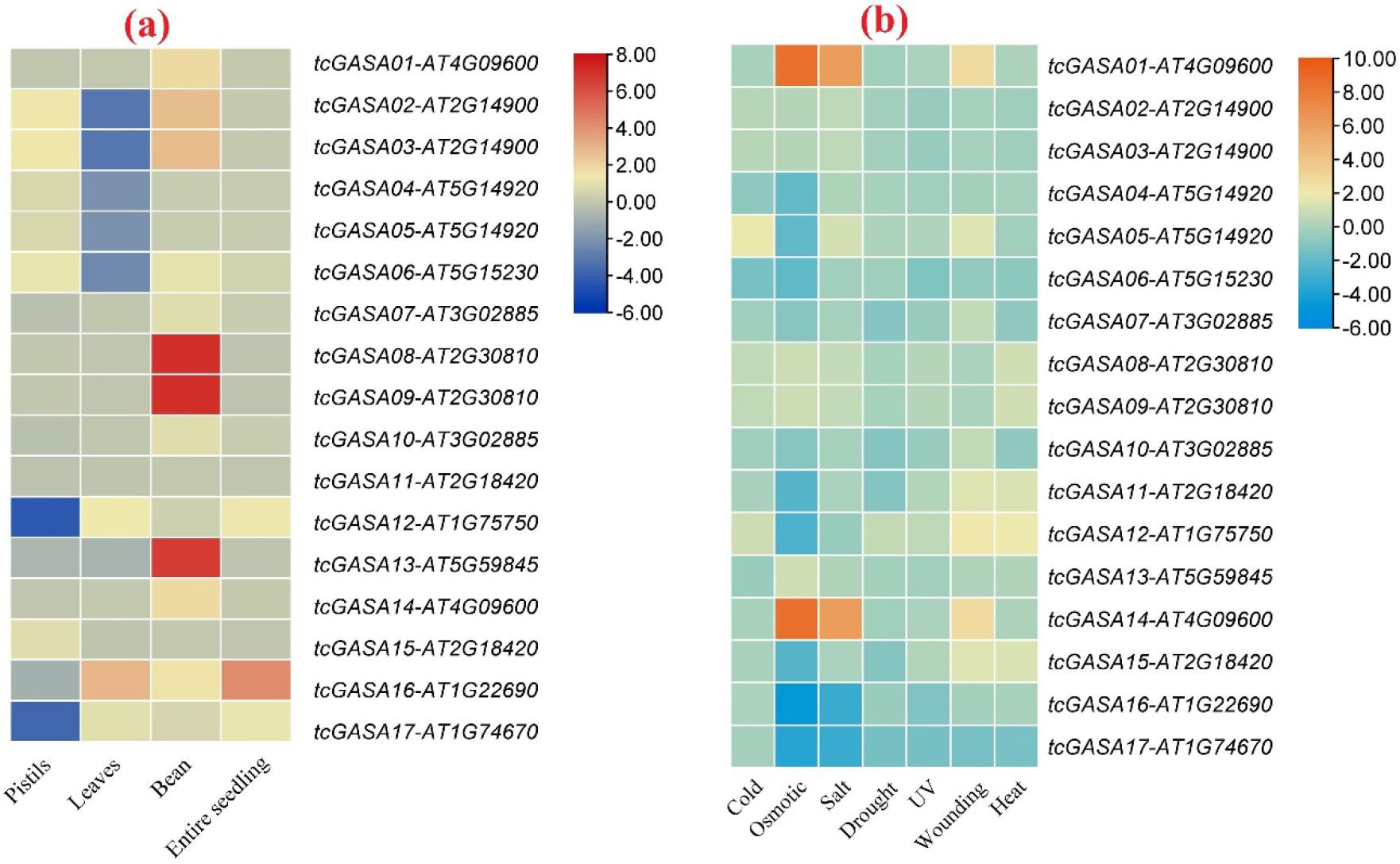
Expression analyses of orthologous *tcGASA* genes of Arabidopsis. (a) Expression in different types of tissues, (b) expression in response to various abiotic stresses.

### 3.10 Expression analyses of *tcGASA*s in abiotic and biotic stresses

We explicated the role of *tcGASAs* under various abiotic stress by analyzing orthologous genes of Arabidopsis (Figure 8b). The orthologous genes of tcGASA01, *tcGASA*05, *tcGASA*12, and *tcGASA*14 showed an upregulation in response to wound healing and the orthologs of *tcGASA*05 were more expressed in response to cold stress. The expression of orthologous *GASA* genes in Arabidopsis was less induced in response to drought and UV stresses. The *tcGASA*01 and *tcGASA*14 were highly upregulated in response to osmotic pressure and salt stress, while *tcGASA*016 and *tcGASA*17 were downregulated.

### 3.11 Expression analyses of *tcGASA*s in biotic stress (*P. megakarya*)

The role of *tcGASAs* against *P. megakarya* was accessed using RNA-seq data of cacao inoculated with *P. megakarya* for 0 hour (0h) 6h, 24h, and 72 h in fungal resistant cultivar Scavina (SCA6) and susceptible cultivar Nanay (NA32) (Figure 9a, b). Four genes *tcGASA*01, *tcGASA*08, *tcGASA*09, *tcGASA*15 were not expressed. The analyses showed differential expression of *tcGASA*s expression under biotic stress of fungus *P. megakarya* in SCA6 and NA32 (Figure 10). The *tcGASA*03, *tcGASA*05, *tcGASA*06, *tcGASA*16, and *tcGASA*17 showed high expression after 24 hours (24h) of treatment and *tcGASA*02, *tcGASA*04, and *tcGASA*13 highly expressed after 72h only in SCA6. This showed their possible role to fungal resistant.

**Figure 9.**
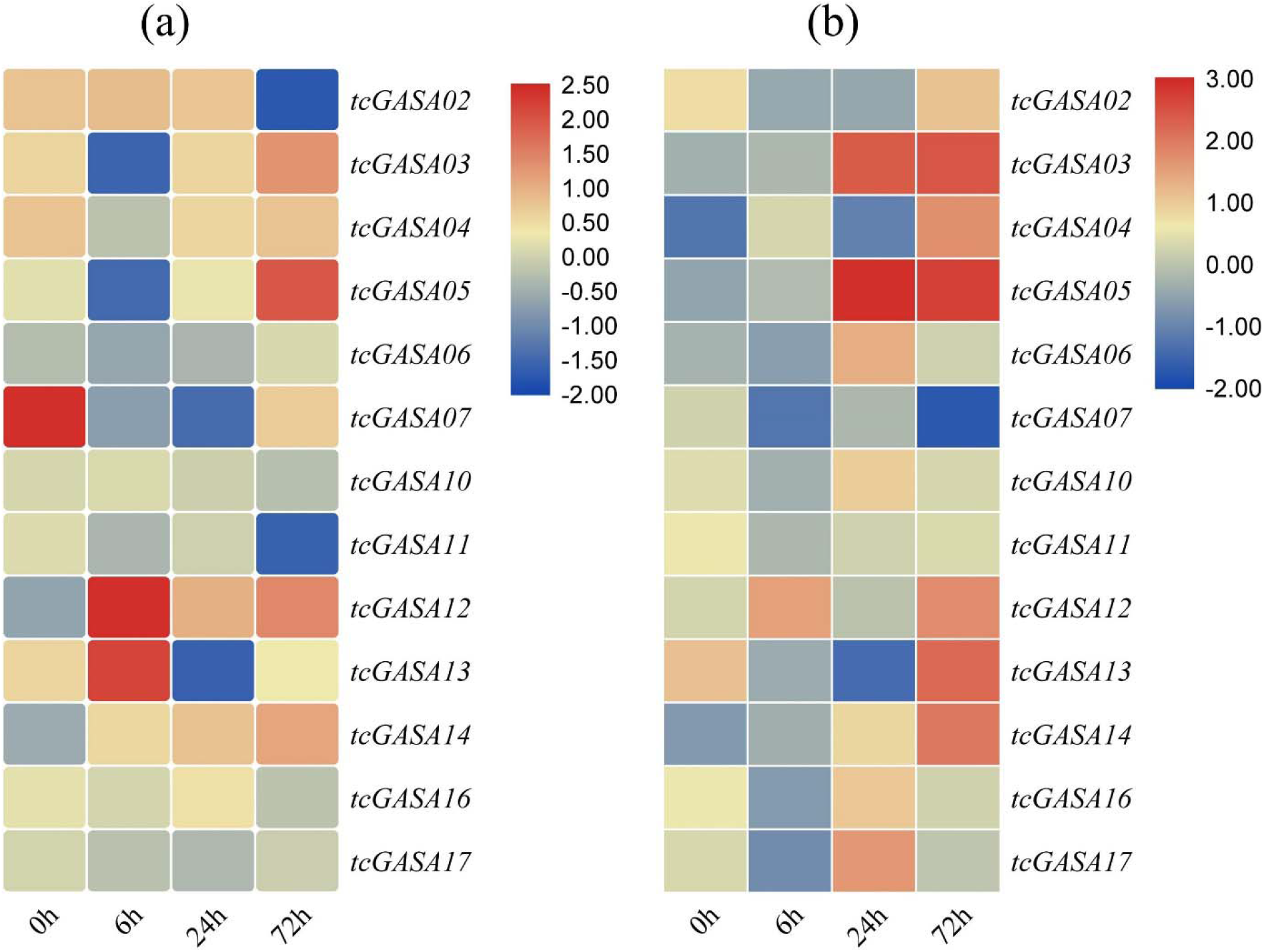
Expression analyses of *tcGASA* genes in cacao plants after inoculation with *P. megakarya* after 0h, 6h, 24h, and 72h. (a) Nanay (NA-32) susceptible cultivar, (b) Scavina (SCA6) tolerant cultivar.

**Figure 10.**
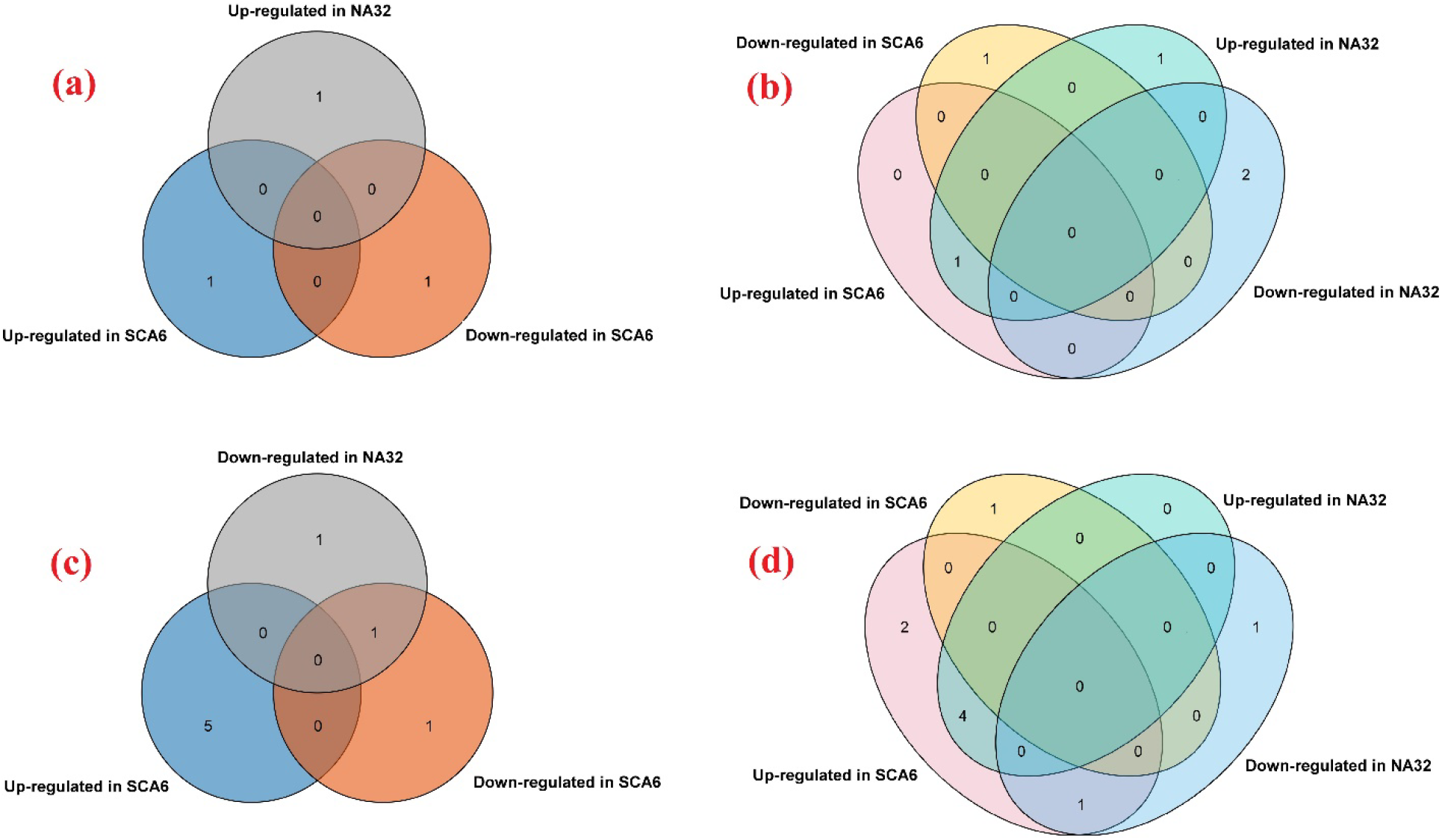
Venn diagram to represent differential expression in susceptible cultivar Nanay (32) and tolerant Cultivar (SCA6) of cacao. (a) 0h (b) 6h (c)24h (d) 72h

## 4. Discussion

In the current study, we identified 17 tcGASAs and analyzed their genomic distributions and chemical properties. RNAseq data analyses explicated their possible roles in bean development (food of cacao), various abiotic stresses, and the biotic stress of *P. megakarya*.

Cacao is an economically important plant and its seeds are used for chocolate, and it is also the main source of income for 40–50 million farmers (Abdullah et al., 2019b; Litz, 2005; Motamayor et al., 2013). Despite this importance, only two gene families, *WRKY* (Dayanne et al., 2017) and *NAC* (Shen et al., 2020), are studied in cacao. The role of the *NAC* family was not explained in abiotic and biotic stresses. The *GASA* family’s role was reported in plant development, function regulation, and biotic and abiotic stresses, as mentioned in the introduction (*vide infra*). Cacao production is also affected by various biotic and abiotic stresses. In the current study, we focused on the GASA gene family in cacao. We identified 17 GASA genes, which were unequally distributed on six chromosomes among ten. The GASA proteins have low molecular weight with conserved GASA and cysteine domains. Our findings are in agreement with previous studies that revealed that GASA genes mostly exist in lower numbers, have low molecular weight, and are unequally distributed on chromosomes within genomes as reported for rice (9; Rezaee et al., 2020), Grapevine (14; Ahmad et al., 2020b), Arabidopsis (15; Fan et al., 2017), and tomato (19; Rezaee et al., 2020). However, a somewhat high number of GASA have also reported, such as in apple (26; Fan et al., 2017) and soybean (37; Ahmad et al., 2019). The number of introns were variable in tcGASA within genes that cluster together in phylogeny. The loss and gain of introns occurs in the course of evolution within protein-coding genes of the plants and is also reported within the GASA of other plant species (Abdullah et al., 2020a; Ahmad et al., 2019, 2020b; Ahmadizadeh et al., 2020; Ding et al., 2020; Heidari et al., 2019).

The tcGASAs were predicted to be alkaline, hydrophilic, and mostly unstable proteins. However, we also detected four stable proteins: tcGASA3, tcGASA11, tcGASA13, and tcGASA15. The stability demonstrates the lifetime of proteins related to cellular enzymatic reactions (Heidari et al., 2019). Hence, these four proteins may play extensive roles in various enzymatic activities. The data of subcellular localization also provides insight into the function of proteins (Zhang and Wang, 2011). Apart from tcGASA8 (which localizes to the plasma membrane), other proteins locate in the extracellular space. Extracellular localization of GASA proteins in several plants has also been reported previously (Ahmad et al., 2019; Rezaee et al., 2020; Zhang et al., 2009). Localization of GASA proteins in the plasma membrane, cytoplasm, and the nucleus has also been described, however (Wang et al., 2009). Variations in subcellular localization may occur due to various factors, such as protein-protein interaction and post-translational modifications (Nahirñak et al., 2016). Post-translational modifications are processes of chemical modifications of proteins and they produce diversity in structure and function, including subcellular localization, protein-protein interaction, and regulating enzyme activity by allosteric phenomena (Duan and Walther, 2015; Nahirñak et al., 2016; Webster and Thomas, 2012). We predicted phosphorylation sites in all tcGASA, ranging in number from 7 to 57. The phosphorylation of proteins is also vital for cell signaling, regulation of various protein mechanisms, and as a substrate for various kinases (Heidari et al., 2020; Nawaz et al., 2019; Silva-Sanchez et al., 2015). Hence, tcGASA may be a good target for various studies to elucidate their complete role. Similarly, glycosylation is also an abundant and varied modification that plays an essential role in the biological and physiological functions of a living organism (Yang et al., 2017). We detected glycosylation sites on the N-terminal of tcGASA10, tcGASA14, and tcGASA16. These tcGASAs may be play significant roles in plant function and regulations.

We performed the phylogenetic analyses of tcGASAs with GASA of 6 other species. We also included *Gossypium raimondii,* a closely related species from the plant family Malvaceae. The phylogenetic analyses distributed GASA proteins into 5 groups and the tcGASAs were also distributed into all five groups. Tc08v2_t01469 (tcGASA16) was the only protein which grouped with Arabidopsis, while other tcGASAs showed sister relationships with GASA of *Gossypium*. The cacao belongs to the basal group, whereas *Gossypium* belongs to the crown group of the family Malvaceae (Abdullah et al., 2019a, 2020c, 2020b), but the close phylogenetic relationships of proteins of these two species support their family-level relationship. The motifs and exon-intron analyses showed variations within proteins that cluster together in phylogeny. This shows that GASA of some groups may have evolved during evolution, which led to variations in motifs and introns in some groups. The same was observed in GASA of other species and has also been reported for other gene families (Ahmad et al., 2019, 2020b; Chattha et al., 2020; Lei et al., 2020). The phylogenetic grouping of GASA was not correlated with the function of the GASA, and we observed contrasting gene expression, including in the same group. These findings show that some other processes are related to the function of proteins instead of their close phylogenetic relationships. A similar observation was reported in *Nicotiana tabacum* L. (Nawaz et al., 2019). However, some studies also proposed that the closely related proteins on a phylogenetic tree have similar function (Nawaz et al., 2014; Nuruzzaman et al., 2010).

The tandem and segmental duplicated genes play an important role in evolution, domestication, functional regulation, and biotic and abiotic stresses (Ahmad et al., 2020a; Li et al., 2020; Liu et al., 2020; Piégu et al., 2020; Song et al., 2020). We determined segmental duplication of nine genes that form four groups as: *tcGASA*2-*tcGASA*3-*tcGASA*13, *tcGASA*8-*tcGASA*9, *tcGASA*14-*tcGASA*15, and *tcGASA10*-*tcGASA17*. These gene pairs were present in the same group within the phylogenetic tree. Similar results were reported in genome-wide analyses of *GASA*s in soybean (Ahmad et al., 2019). A previous study also proposed that segmentally duplicated genes also showed similar functions and stable expressions. (Ahmad et al., 2019; Faraji et al., 2020). In the current study, each pair of segmentally duplicated genes did not agree with the previous finding. The pairs *tcGASA*8-*tcGASA*9 and *tcGASA*14-*tcGASA*15 showed similarities in gene expression in different tissues. The *tcGASA*2-*tcGASA*3-*tcGASA*13 showed a similar expression in the entire seedling and bean but showed differential expression in pistils and leaves. In biotic stress of fungus (*P. megakarya*), *tcGASA*2-*tcGASA*3-*tcGASA*13, and *tcGASA*8-*tcGASA*9, *tcGASA10*-*tcGASA17* showed similar expression, while *tcGASA*14-*tcGASA*15 and showed different expression as *tcGASA*14 showed high expression whereas *tcGASA*15 was found unexpressed. These findings suggest that segmentally duplicated pairs may also perform different functions. Hence, functional analyses of each segmentally duplicated gene can provide authentic information about their roles in various physiological and biochemical processes. Moreover, our study, along with the previous report (Ahmad et al., 2019), suggests that genes within the same groups have more chances of segmental duplication events among them. The Ka/Ks < 1 indicates that purifying selection pressure exists on the *GASA* genes after duplication, as reported previously (Ahmad et al., 2020b). We also observed mostly purifying selection pressure on *tcGASA*s, including duplicating genes (Table S1).

The cis-acting regulatory elements involved in transcription of regulation genes are induced through independent signal transduction pathways under biotic and abiotic stresses (Ahmadizadeh and Heidari, 2014; Nawaz et al., 2019). We observed several key cis-regulating elements in response to light, hormones, stresses, and growth in the promoter site of *tcGASAs*. The cis-regulating elements for drought, anerobic induction, low-temperature, and plant defense were also evident. The existence of diverse cis-regulating elements in promoter regions indicates their roles in the regulation of the *tcGASAs* and different pathways of cacao. Further study can shed light on the roles of these cis-regulating elements.

Different strategies are used to develop cacao cultivars that produce food in high quantities and are resistant to abiotic and biotic stresses (Bridgemohan and Mohammed, 2019). Here, we studied the role of *GASA* genes in cacao development and in protection against biotic and abiotic stresses. Tissue-specific expression was observed for *tcGASAs*, which showed their role in the development and functional regulation of cacao. Up to eight *tcGASAs* were highly expressed in the cacao bean, which is the food part used for making chocolate (Abdullah et al., 2020d). This high expression may reveal the conserved function of these genes in the development of the bean. The role of *tcGASA*s was also stated in the development of grapevine seeds (Ahmad et al., 2020b). Further data based on cloning can provide new insight into their roles in bean development. Expression analyses of the orthologous genes of Arabidopsis also indicated the role of *GASA* genes in various abiotic stresses, including drought, which significantly affects the growth of cacao (Medina and Laliberte, 2017). Hence, these genes may also be important to produce drought-resistant cultivars. Expression analyses also provide insight into the gene’s function in response to biotic stresses (Nawaz et al., 2019). The black rod disease of genus *Phytophthora* caused up to 20–25% loss (700,000 metric tons) to the world cacao production annually. In some regions, the *Phytophthora* caused losses of 30–90% of the crops (Adeniyi, 2019). Here, we explored the function of *tcGASA*s based on RNAseq data against black rod causing pathogen *P. megakarya* and observed highly expressed *tcGASA* genes in plants inoculated at 24h and 72h in the tolerant cultivar SCA6 as compared to susceptible cultivar NA32. This data indicates that *tcGASAs* respond to fungus. Hence, the complete characterization of these upregulated genes can provide target genes for the development of resistant cultivars to the disease of genus *Phytophthora* to enhance the production of cacao for the welfare of not only farmers involved in cacao cultivation, but also for the welfare of all humanity to provide high-quality delicious chocolate with quality nutrition.

In conclusion, our study provides new insight into the identification, characterization, and expression of the *GASA* genes in the *Theobroma cacao* plant. Expression analyses revealed the role of the *GASA* genes in seed development. We also determined that *GASA* genes provide resistance against the fungus *Phytophthora megakarya,* which causes significant losses to cacao production each year. Our study provides a base for the generation of cultivars that are resistant to fungus of the genus *Phytophthora*.

## Data availability

The public data set is analyzed in the manuscript. All the analyzed data are available in the main manuscript or as supplementary materials.

